## Supplementary material for "Effects of route of administration on oxytocin-induced changes in regional cerebral blood flow in humans"

Martins et al.

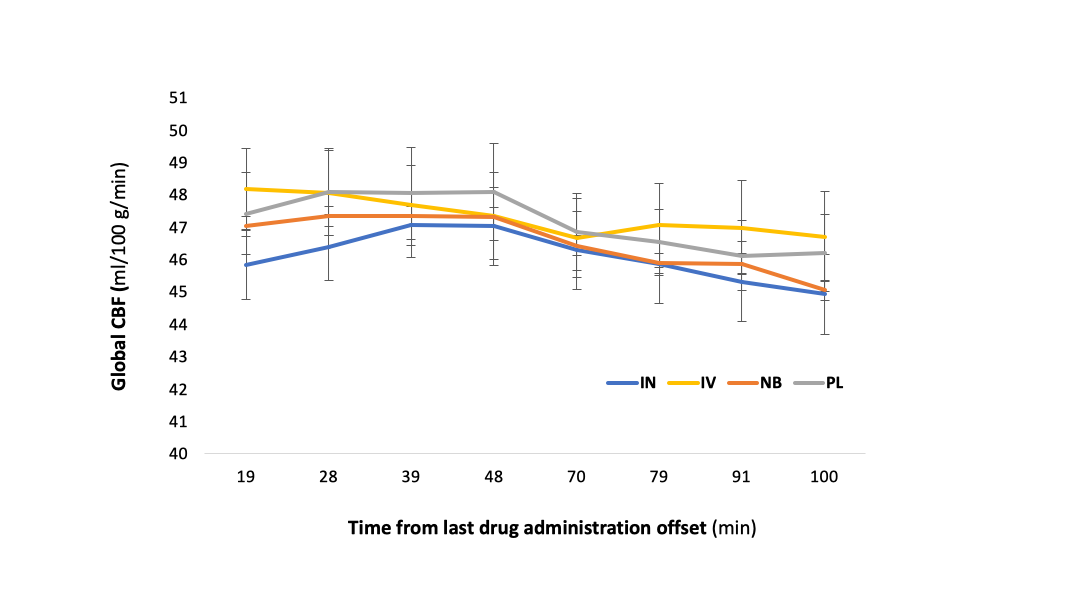

**Supplementary Figure 1** – **Global cerebral blood flow across time.** We tested for treatment, time interval and treatment x time-interval effects on global cerebral blood flow (CBF) signal in a 2-way repeated measures analysis of variance. Data are presented as mean ± 1 standard error of the mean (SEM). Statistical significance was set to p<0.05 (two-tailed). Error bars correspond to SEM. IN – Spray; IV – Intravenous; NB – Nebulizer; PL – Placebo.

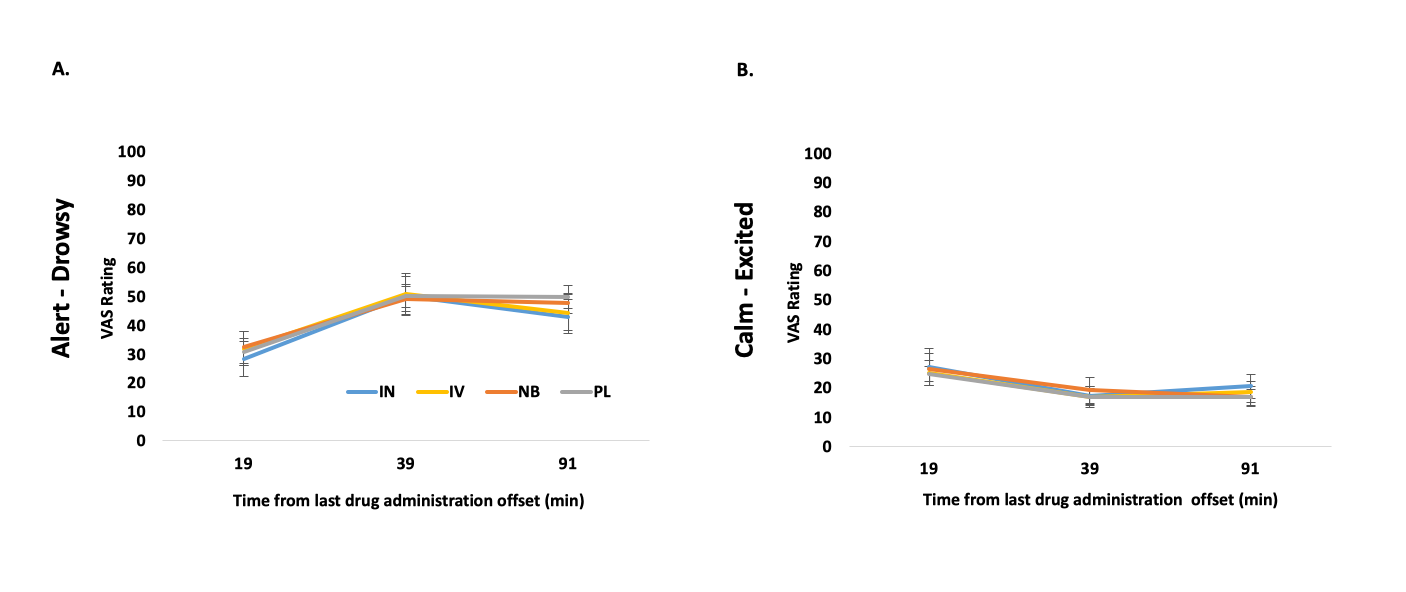

**Supplementary Figure 2** – **Subjective ratings of alertness and excitement** We tested for treatment, time interval and treatment x time-interval effects on subjective ratings of alertness and excitement, collected at 3 different time-points during the scanning period, in a repeated measures analysis of variance. In panels A and B, we present the results for alertness and excitement, respectively. Anchors - Alertness: 0 – Alert; 100 – Drowsy; Excitement: 0 – Calm; 100 – Excited. Data are presented as mean ± 1 standard error of the mean (SEM). Statistical significance was set to p<0.05 (two-tailed). Error bars correspond to SEM. IN – Spray; IV – Intravenous; NB – Nebulizer; PL – Placebo.

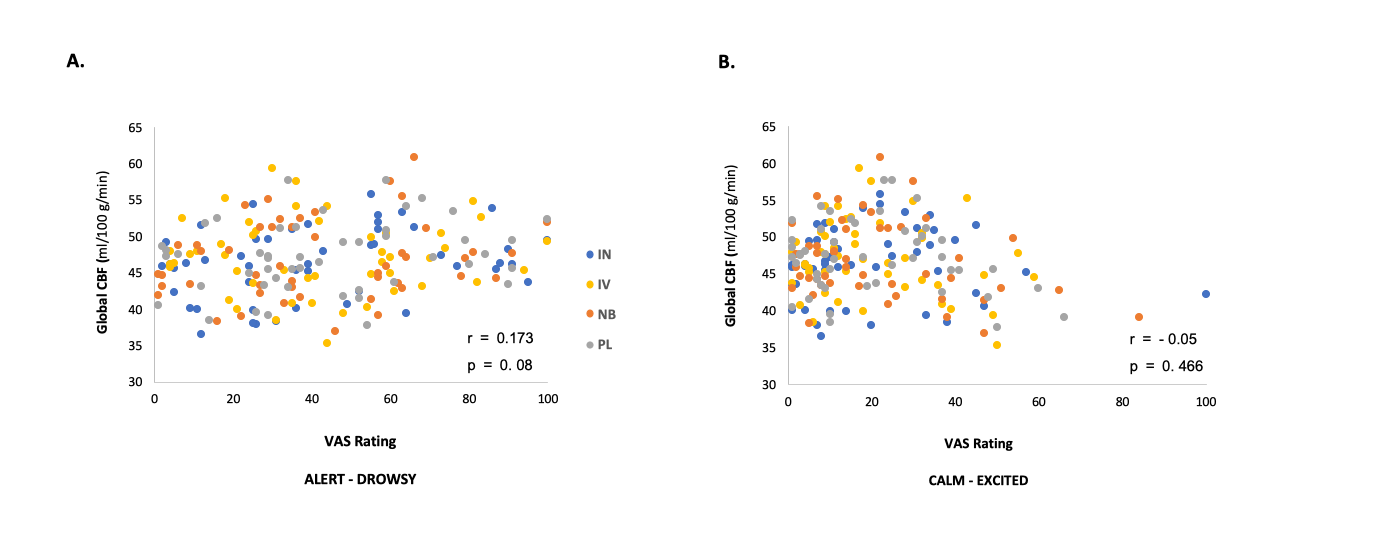

**Supplementary Figure 3** – **Correlation between global cerebral blood flow and subjective ratings.** To investigate if the changes in the subjective ratings we identified are associated with changes in global cerebral blood flow (CBF) across time, we used within-group pooled Pearson correlation coefficients (r) to examine self-ratings of alertness and excitement could predict global CBF over time. In panel A, we present Global CBF vs Alertness; and in panel B, Global CBF vs Excitement. Statistical significance was set to p<0.05 (two-tailed). IN – Spray; IV – Intravenous; NB – Nebulizer; PL – Placebo.

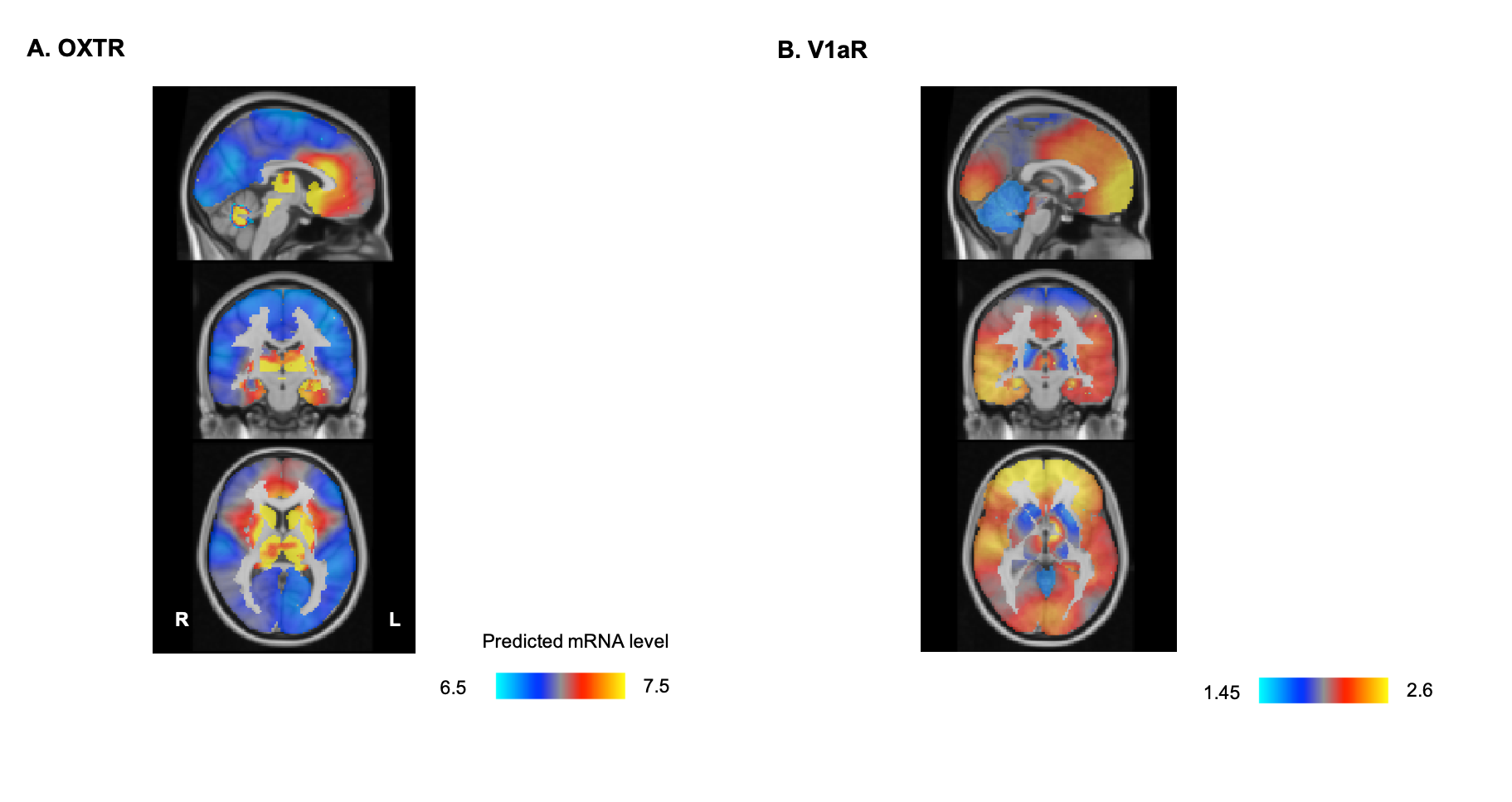

**Supplementary Figure 4** – **Human whole-brain distribution of OT-related receptors.** Panel A refer to OXTR and Panel B to V1aR. Scale refers to spatial whole-brain distribution of the predicted mRNA expression intensity as determined by Gryglewski et al.^1^ using the Allen Brain Atlas data; these maps were generated by applying a variogram model in Gaussian process regression allowing to infer gene expression with high resolution and across the whole brain; the original maps can be freely downloaded from http://www.meduniwien.ac.at/neuroimaging/mRNA.html)

**Supplementary Table 1 – Effects of treatment, time-interval and treatment x time-interval on global cerebral blood flow and subjective ratings.** Detailed statistics for each effect. The effects of treatment, time-interval and treatment x time-interval were tested using repeated measures 2-way ANOVA. Statistical significance was set to p<0.05 (two-tailed). * highlights statistically significant effects; CBF – Cerebral blood flow.

|  | Main effect of treatment | | | Main effect of time-interval | | | Treatment x Time-interval interaction | | |
| --- | --- | --- | --- | --- | --- | --- | --- | --- | --- |
|  | F | p | η^2^_p_ | F | p | η^2^_p_ | F | p | η^2^_p_ |
| Global CBF | 1.436 | 0.245 | 0.062 | 6.597 | <0.0001* | 0.793 | 0.661 | 0.870 | 0.058 |
| Alertness | 0.301 | 0.825 | 0.091 | 38.200 | <0.0001* | 0.793 | 0.715 | 0.639 | 0.326 |
| Excitement | 0.424 | 0.737 | 0.059 | 6.759 | 0.0038* | 0.486 | 0.834 | 0.834 | 0.142 |

**Supplementary Table 2** **– Pharmacokinetics analysis**. We present a summary of three parameters describing the pharmacokinetics of each treatment administration method. We present the maximum plasmatic concentration achieved after dosing (C_max_), the area under the curve (AUC) reflecting the magnitude of the variations of plasmatic oxytocin post-dosing and the absolute bioavailability of oxytocin absorbed to the plasma, calculated as the dose-corrected AUC of each intranasal administration divided by the dose-corrected AUC of the intravenous administration. Data are presented as mean (1 standard error of the mean).

| Method | C_max_ (pg.ml^-1^) | AUC (pg.ml^-1^.h^-1^) | Absolute bioavailability (%) |
| --- | --- | --- | --- |
| Intravenous | 64.314 (4.702) | 284.473 (26.538) | NA |
| Spray | 15.790 (1.775) | 159.459 (18.931) | 16.390 (2.889) |
| Nebulizer | 16.317 (1.767) | 115.022 (7.165) | 12.160 (1.686) |

**Supplementary Table 3 – Spearman correlations between drug-induced changes in rCBF and variations of the concentration of oxytocin in the plasma post-dosing.** We present values of the spearman correlation coefficient (r), with respective 95% confidence intervals (CI) for all clusters identified as significant in the comparisons drug vs placebo for the three treatment methods of administration. Statistical significance was set to p<0.05 (two-tailed) and is highlighted with a star symbol.

| Cluster | r (CI 95%) | p |
| --- | --- | --- |
| Spray vs Placebo | | |
| 1 | 0.050 (-0.469 to 0.543) | 0.857 |
| 2 | - 0.502 (-0.805 to 0.006) | 0.049 |
| 3 | 0.477 (-0.041 to 0.793) | 0.064 |
| 4 | 0.102 (-0.427 to 0.580) | 0.681 |
| 5 | 0.111 (-0.419 to 0.586) | 0.680 |
| 6 | - 0.52 (-0.813 to -0.017) | 0.041 |
| 7 | - 0.213 (-0.421 to 0.134) | 0.092 |
| Intravenous vs Placebo | | |
| 1 | - 0.544 (-0.824 to -0.050) | 0.031 |
| 2 | - 0.567 (-0.834 to -0.084) | 0.024 |
| Nebulizer vs Placebo | | |
| 1 | - 0.268 (-0.683 to 0.278) | 0.315 |
| 2 | 0.035 (-0.481 to 0.534) | 0.899 |
| 3 | 0.050 (-0.469 to 0.543) | 0.857 |
| 4 | 0.137 (-0.835 to -0.321) | 0.457 |
| 5 | 0.277 (-0.269 to 0.688) | 0.299 |

**Supplementary Table 4 – Percentage of observations satisfying quality control criteria for the pulse plethysmography analysis.** Following current recommendations for heart rate data processing and analysis, if more than 5% of the beats required correction, then we decided to exclude these periods of observation. In some rare cases where artefacts could not be corrected, we only included the data if at least 5 min of acquisition free of artefacts could be analysed. In this table, we present the percentage of total data available satisfying our quality control criteria for each treatment method of administration and time-interval, so the reader can appreciate the amount of data effectively used in the final analysis.

|  | Time from last drug administration offset (minutes) | | | | | | | |
| --- | --- | --- | --- | --- | --- | --- | --- | --- |
| Method | 15-23 | 24-32 | 35-43 | 44-52 | 66-74 | 75-83 | 87-95 | 96-104 |
| Spray | 100% | 93.75% | 87.5% | 100% | 100% | 87.5% | 87.5% | 87.5% |
| Intravenous | 100% | 100% | 100% | 100% | 100% | 100% | 93.75% | 87.5% |
| Nebulizer | 100% | 93.75% | 100% | 100% | 100% | 93.75% | 93.75% | 100% |
| Placebo | 87.5% | 87.5% | 87.5% | 87.5% | 87.5% | 87.5% | 81.25% | 87.5% |

**Supplementary Table 5** – **Oxytocin did not affect heart rate or heart rate variability.** We determined time-interval, treatment, and treatment x time-interval effects on a set of time domain (heart-rate (HR) and the root mean square of the successive differences (RMSSD), frequency domain (low (LF) and high (HF) frequencies spectral powers and high/low frequency spectral power ratio (HF/LF)) and non-linear (Approximate entropy (ApEn), the SD1 and SD2 lines from the Poincare Plot and the detrended fluctuation scaling exponents DFAα1 and DFAα2) analysis measures of heart rate variability. We used a repeated measures two-way analysis of variance, using subjects, treatment and time as factors, and the Greenhouse-Geisser correction against violations of sphericity. Statistical significance was set to p<0.05. We contained the false-discovery rate (FDR) at α=0.05 using the Benjamini-Hochberg procedure. Original p values (two-tailed) are reported alongside with values obtained after accounting for FDR.

|  | | Main effect of treatment | | | | | Main effect of time | | | | | Interaction Treatment x time | | | |
| --- | --- | --- | --- | --- | --- | --- | --- | --- | --- | --- | --- | --- | --- | --- | --- |
|  |  | F | P (unadjusted) | P (FDR) | η^2^_p_ | F | | P (unadjusted) | P (FDR) | η^2^_p_ | F | | P (unadjusted) | P (FDR) | η^2^_p_ |
| Time-domain | HR | 0.093 | 0.832 | 8.320 | 0.015 | 2.359 | | 0.14 | 0.156 | 0.282 | 0.912 | | 0.465 | 2.325 | 0.132 |
|  | RRMSD | 1.605 | 0.223 | 1.115 | 0.037 | 1.511 | | 0.233 | 0.233 | 0.304 | 0.485 | | 0.722 | 0.903 | 0.131 |
| Frequency-domain | HF | 0.314 | 0.682 | 0.853 | 0.050 | 3.627 | | 0.040 | 0.067 | 0.377 | 0.777 | | 0.535 | 2.325 | 0.115 |
|  | LF | 0.442 | 0.668 | 0.954 | 0.069 | 5.776 | | 0.005 | 0.050 | 0.490 | 0.660 | | 0.635 | 0.907 | 0.099 |
|  | HF/LF | 1.034 | 0.396 | 0.792 | 0.147 | 3.804 | | 0.029 | 0.097 | 0.388 | 0.370 | | 0.847 | 0.847 | 0.058 |
| Nonlinear | SD1 | 0.231 | 0.775 | 0.861 | 0.037 | 2.626 | | 0.097 | 0.139 | 0.304 | 0.906 | | 0.465 | 1.550 | 0.131 |
|  | SD2 | 0.050 | 0.985 | 0.985 | 0.008 | 4.078 | | 0.029 | 0.073 | 0.405 | 0.763 | | 0.556 | 0.927 | 0.113 |
|  | ApEN | 0.715 | 0.498 | 0.830 | 0.107 | 1.988 | | 0.132 | 0.165 | 0.249 | 0.964 | | 0.445 | 4.450 | 0.138 |
|  | DFA alfa1 | 1.711 | 0.218 | 0.727 | 0.222 | 4.800 | | 0.017 | 0.085 | 0.444 | 0.382 | | 0.844 | 0.938 | 0.060 |
|  | DFA alfa2 | 1.267 | 0.315 | 0.788 | 0.174 | 3.675 | | 0.033 | 0.066 | 0.380 | 0.814 | | 0.533 | 1.333 | 0.119 |
